## Supplementary material for "Aspartate reduces liver inflammation and fibrosis by suppressing the NLRP3 inflammasome pathway via upregulating NS3TP1 expression": author cover letter

Running title: Aspartate prevents liver fibrosis

The author information is as follows:

Li Zhou, PhD, Department of Infectious Disease, China-Japan Friendship Hospital, 100029, Beijing, China;

Jing Zhao, PhD, Institute of Infectious Diseases, Beijing Ditan Hospital, Capital Medical University, Beijing 100015, China;

Ming Han, PhD, Institute of Infectious Diseases, Beijing Ditan Hospital, Capital Medical University, Beijing 100015, China;

Shunai Liu, MD, MS, Institute of Infectious Diseases, Beijing Ditan Hospital, Capital Medical University, Beijing 100015, China;

Xiaoxue Yuan, PhD, Institute of Infectious Diseases, Beijing Ditan Hospital, Capital Medical University, Beijing 100015, China;

Anlin Ma, MS, Chief physician, Department of Infectious Disease China-Japan Friendship Hospital, 100029, Beijing, China;.

Song Yang, MD, PhD, Institute of Infectious Diseases, Beijing Ditan Hospital, Capital Medical University, Beijing 100015, China;

Yilan Zeng, MD, Chengdu Public Health Clinical Medical Center,

Jun Cheng, MD, PhD, EMBA, Institute of Infectious Diseases, Beijing Ditan Hospital, Capital Medical University, Beijing 100015, China;

Corresponding author: Jun Cheng, MD, PhD, EMBA, Institute of Infectious Diseases, Beijing Ditan Hospital, Capital Medical University, NO.8 East Jingshun Street,Chaoyang, Beijing 100015, China. Phone: +86 10 8432 2006, Fax: +86 10 8439 7196,. Yilan Zeng, MD, Chengdu Public Health Clinical Medical Center, 377 Jingming Rd, Jinjiang District, Chengdu City, Sichuan Province 610061, China;
