## Supplementary material for "Aspartate reduces liver inflammation and fibrosis by suppressing the NLRP3 inflammasome pathway via upregulating NS3TP1 expression": letter to the editor

Dear editors,

We specialize in the pathogenesis of liver fibrosis including cell proliferation, apoptosis, steatosis, autophagy and inflammation. Previous studies have suggested that aspartate can act on liver Kupffer cells, inhibit NLRP3 inflammatory bodies and improve liver inflammation in acute hepatitis. In this study we sought to explore the roles of aspartate in chronic liver injury, human liver fibrosis. We found aspartate plays as a compelling inflammation inhibitor, which suppresses liver fibrosis by targeting NLRP3 pathway. Four points are highlighted as follows.

1. We confirmed that NLRP3 inflammasome signaling pathways induced cell inflammation and activated hepatic stellate cells, which is a potential strategy in both acute and chronic hepatitis.
2. We showed that aspartate inhibits fibrogenic activation of hepatic stellate cells and prevents liver fibrosis *in vivo* and *in vitro* via NF-κB/NLRP3 infammasome signaling pathways.
3. We illustrated that aspartate inhibits cell proliferation and induces apoptosis and lipid accumulation, which have been shown to be potential factors in liver fibrosis.
4. We demonstrated that NS3TP1 reduced liver fibrosis in LX-2 cells via regulating NF-κB/NLRP3 signaling pathways.

This work was supported by the National Natural Science Foundation of China, and the National Key R&D Program of China. The authors claim that the funding organizations are public institutions and have no role in the design and conduct of this study. No portion of the manuscript has been published or is under consideration for publication elsewhere. The study complies with current ethical considerations.

Thank you for your consideration.

Yours sincerely,

Li Zhou, MD, PhD.

Corresponding author:

Yilan Zeng, MD,

Jun Cheng, MD, PhD, Prof.
